# Neural basis for adaptive motor behavior during car driving

**DOI:** 10.1101/2021.10.03.462866

**Authors:** Ryu Ohata, Kenji Ogawa, Hiroshi Imamizu

**Affiliations:** Department of Psychology, Graduate School of Humanities and Sociology, The University of Tokyo, Hongo 7-3-1, Bunkyo-ku, Tokyo 113-0033, Japan; Department of Psychology, Graduate School of Letters, Hokkaido University, Kita 10, Nishi 7, Kita-ku, Sapporo 060-0810, Japan; Cognitive Mechanisms Laboratories, Advanced Telecommunications Research Institute International (ATR), Keihanna Science City, Kyoto 619-0288, Japan; Research into Artifacts, Center for Engineering, The University of Tokyo, Hongo 7-3-1, Bunkyo-ku, Tokyo 113-0033, Japan

**Author notes:** Correspondence and requests for materials should be addressed to H.I or R.O.

**Keywords:** car driving, motor control, internal model, frontoparietal network, default mode network, salience network

## Abstract

Car driving is supported by motor skills trained through continuous daily practice. One of the skills unique to expert drivers is the ability to detect abrupt changes in the driving environment and then quickly adapt their operation mode to the changes. Previous functional neuroimaging studies on motor control investigated the mechanisms underlying behaviors adaptive to changes in control properties of simple experimental devices such as a computer mouse or a joystick. The switching of multiple internal models mainly engages adaptive behaviors and underlies the interplay between the cerebellum and frontoparietal network (FPN) regions as the neural process. However, it remains unclear whether the neural mechanisms identified through an experimental paradigm using such simple devices also underlie practical driving behaviors. In the current study, we measure functional magnetic resonance imaging (fMRI) activities while participants control a realistic driving simulator inside the MRI scanner. Here, the accelerator sensitivity of a virtual car is abruptly changed, requiring participants to respond to this change as quickly as possible. We first compare brain activities before and after the sensitivity change. As a result, sensorimotor areas, including the left cerebellum, increase their activities after the sensitivity change. Moreover, after the change, activity significantly increases in the inferior parietal lobe and dorsolateral prefrontal cortex, parts of the FPN regions. By contrast, the posterior cingulate cortex, a part of the default mode network, deactivates after the sensitivity change. Our results suggest that the neural bases found in previous experiments using the simpler devices can serve as the foundation of adaptive car driving. At the same time, this study also highlights the unique contribution of non-motor-related regions to addressing the high cognitive demands of driving.

## 1. Introduction

A car is one of the most advanced devices humans operate in daily life. People need to accumulate driving experience to acquire practical driving skill. Expert drivers, such as racing drivers, are characterized by the ability to rapidly detect changes in the driving environment and flexibly switch their driving modes in response to those changes. For example, expert drivers can maintain their car ‘s movement at a constant speed regardless of running uphill or downhill, and they can keep their driving stable after their car suddenly encounters a slippery road surface. Such adaptive driving behaviors are supported by sophisticated computation in the brain, although experts achieve adaptive behaviors without any difficulty. Knowledge of human motor control, which has been extensively studied in the fields of psychology and neuroscience, could play an important role in understanding the neural mechanism underlying skillful driving behaviors (Lappi, 2015). However, only a limited number of studies have investigated the neural basis for driving from the perspective of human motor control.

A neural system that may be essential to adaptive driving behaviors is internal models, which mimick the input–output properties of our body or controlled devices (Wolpert et al., 1995; Wolpert and Kawato, 1998; Kawato, 1999). The brain maintains not a single but multiple internal models employed for different environments or properties of control devices. Previous neuroimaging studies suggest that the motor-related regions, especially in the cerebellum, maintain multiple internal models in a modular manner (Imamizu et al., 2003; Krakauer et al., 2004; Girgenrath et al., 2008). Imamizu et al. (2003) demonstrated that different types of sensorimotor perturbations (velocity vs. rotation control of a computer mouse) were organized with spatially segregated patterns in the cerebellum. The neural mechanism that allows flexible switching of multiple internal models has also been studied (Imamizu et al., 2004; Imamizu and Kawato, 2008). The frontoparietal network (FPN) regions play a key role in internal model switching through interacting with the cerebellar cortex. The roles of the FPN regions are different; the superior parietal lobe (SPL) is involved in switching depending on predictive contextual information, while the dorsolateral prefrontal cortex (DLPFC) and inferior parietal lobe (IPL) contribute to switching driven by sensorimotor feedback (Imamizu and Kawato, 2008). Thus, the acquisition and switching of multiple internal models is fundamental to adaptive motor control. Since driving skills are mainly supported by motor control functions, the previous literature on human motor control provides clues to understanding the neural mechanisms involved in adaptive driving behavior. However, previous findings were based on well-controlled experimental paradigms using simple devices such as a computer mouse or a joystick. It remains unknown whether the neural bases recruited in controlling such simple devices also support the skills needed for practical car driving.

In the current study, we investigated the brain regions recruited in car driving using functional magnetic resonance imaging (fMRI). Participants drove a virtual car on a simulated circuit course by controlling custom-made devices inside the MRI scanner (Fig. 1A). The ratio of acceleration relative to a stepping-in amount of an accelerator pedal, defined as accelerator sensitivity, abruptly changed in the middle of running over a straight section of the course. Participants were required to respond to the change quickly while maintaining stable driving. We explored activation/deactivation in response to the change in the level of accelerator sensitivity. We also performed multivoxel pattern analysis (MVPA) to examine the cerebellar activity patterns involved in switching driving behavior in response to different levels of accelerator sensitivity.

**Figure 1.**
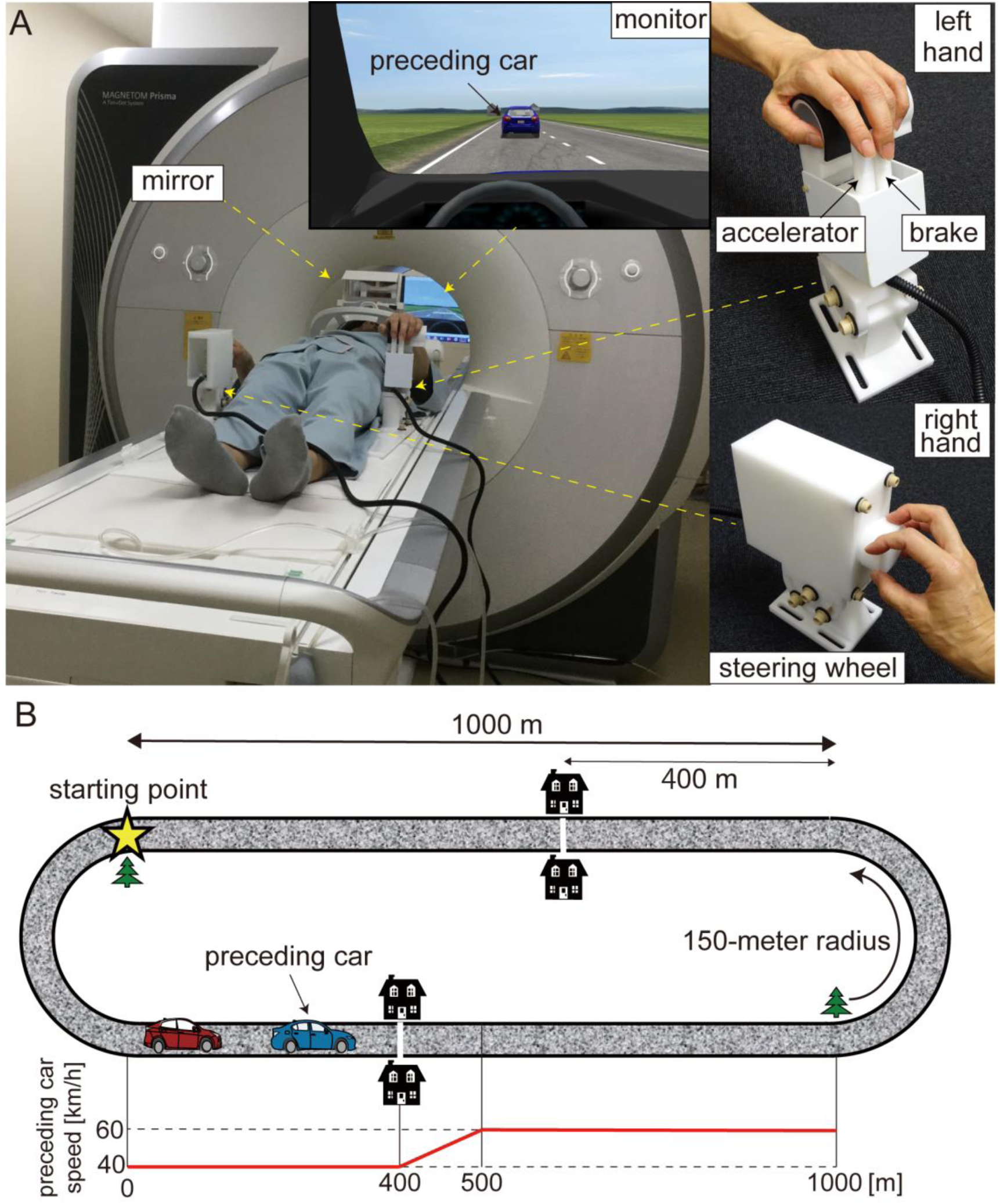
(A) Experimental setup. Participants drove a virtual car while lying on the bed of an MRI scanner. The virtual car was controlled with MRI-compatible customized devices. The two levers manipulated with the index and middle fingers of the left hand served as brake and accelerator pedals (upper right) while the knob controlled with the right hand functioned as a steering wheel (lower right). Participants viewed a monitor via a mirror. (B) Driving course. The driving circuit consisted of 1,000-m straight sections and 150-m radius curves. The star denotes the starting point. The preceding car runs at 40 km/h for the first 400 m of the straight section and accelerates from 40 to 60 km/h for the next 100 m. The white line and two houses were displayed to inform participants of the acceleration point of the preceding car. The bottom panel shows the trajectory of the preceding car’s speed. The trajectory is aligned to the beginning of the straight section (0 m).

## Materials and methods

### Participants

Twenty-five healthy volunteers (three females) with a mean age of 21.9 years (19–33) participated in our experiment. We recruited volunteers who had obtained a driving license. The mean licensed period was 2.7 years (0.4–15). We collected data of 10 participants from the University of Tokyo and 15 participants from Hokkaido University. We excluded the data of five participants from analysis due to large head motion during the fMRI scan (for details, see below: *Preprocessing of MRI data*). We eventually analyzed the data of 20 participants (two females) with a mean age of 22.2 years. Written informed consent was obtained from all volunteers in accordance with the latest version of the Declaration of Helsinki. The experimental protocol was approved by the ethics committee at the University of Tokyo.

### Experimental task

Participants drove a virtual car using a driving simulator (CarSim, Virtual Mechanics, Japan) by controlling a custom-made MRI-compatible control device (Leading-Edge Research and Development Accelerator, Inc., Japan; Fig 1A). They manipulated two levers serving as brake and accelerator pedals with their left hand while controlling a knob serving as a steering wheel with their right hand. We customized program codes with MATLAB and Simulink (Mathworks, USA) to set up the ratio of acceleration/deceleration to a stepping-in amount of the brake and accelerator pedals and that of steering angle to the degree of turning the knob.

A virtual car ran on a circuit course consisting of 1,000-m straight sections and 150-m radius curves (Fig. 1B). Participants were instructed to drive their car while following a preceding car during the task. They were required to keep their car at a 25-m distance from the preceding car as precisely as possible through the straight sections. Note that participants learned how to keep the target distance during the five-day practice session before the fMRI experiment (for details, see below). Although following the preceding car was required, they were not required to attend to the distance between the two cars while driving through the curves. At the start of each run, the participant’s car was placed at the beginning of a curve (star in Fig. 1B). The preceding car ran at 40 km/h for the first 400 m of the straight section and accelerated from 40 to 60 km/h for the next 100 m. A white line across the road and two houses were located in the display 400 m from the beginning of the straight section to inform participants of the timing of the acceleration. Then, the preceding car ran on the remaining 500 -m straight section at 60 km/h (speed trajectory of preceding car is shown in the bottom panel of Fig. 1B, see also Supplementary Movie). We positioned a tree as a landmark of the end of the straight section. The preceding car decelerated from 60 to 40 km/h while going around a curve. We defined the drive through a straight section as one trial. Participants were asked to drive their car four laps in a run, that is, a run consisted of eight trials.

We defined “accelerator sensitivity” as the ratio of the acceleration of a virtual car to the stepping-in amount of an accelerator pedal. We established two levels of accelerator sensitivity (high and low sensitivity), and they abruptly changed from one to the other in the middle of driving on the straight section. Specifically, they changed when the participant’s car crossed the 500-m point from the beginning of the straight section. The high-sensitivity level was three times more sensitive than the low-sensitivity level. This means that a car at high sensitivity produced three times greater acceleration than one at low sensitivity with the same amount of stepping pressure on the accelerator pedal. The level of accelerator sensitivity changed in half of the trials, while not in the remainder of the trials. Participants were informed of the initial accelerator sensitivity at the beginning of each run to reduce the difficulty of driving through the first curve.

### Experiment schedule

We asked each participant to conduct a five-day practice, held outside an MRI scanner, and a two-day MRI session. Note that the practice and MRI sessions were not always conducted on consecutive days. One participant conducted MRI experiments for three days due to his limited schedule. In the practice session, participants drove their car with either low or high sensitivity during an entire run (i.e., without changes in accelerator sensitivity). They practiced driving a car with the two levels of sensitivity in eight runs each throughout the five-day practice (i.e., 16 runs in total). We instructed them to keep a 25-m distance from the preceding car while driving. A white line was displayed 25 m behind the preceding car as guidance of the target distance in the first eight runs of the practice session. In addition to the 16-run practice, participants conducted the same task as the MRI experiment, in which the accelerator sensitivity changed in the middle of driving on a straight section, for two runs at the end of the practice session. The MRI session consisted of five runs per day, and thus we conducted ten runs over two days. Note that three participants (out of 20) performed nine runs in total due to their limited schedule during the MRI scanning.

### MRI data acquisition

A 3 Tesla Magnetom Prisma scanner (Siemens, Germany) with a 64-channel head coil was used to acquire T2*-weighted echo-planar images (EPI). We acquired 435 volumes in each run with a gradient echo EPI sequence under the following scanning parameters: repetition time (TR), 2000 ms; echo time (TE), 30 ms; flip angle (FA), 80°; field of view (FOV), 192×192 mm; matrix, 64×64; 32 axial slices; and thickness, 4 mm with a 1-mm gap. T1-weighted (T1w) magnetization-prepared rapid acquisition gradient-echo (MP-RAGE) fine-structural images were obtained with 1×1×1-mm resolution with a gradient echo sequence (repetition time, 2,250 ms; echo time, 2.98 ms; inversion time (TI), 900 ms; flip angle, 9°; FOV, 256×256; 192 axial slices; and slice thickness, 1 mm without gap). Although we collected MRI data at both the University of Tokyo and Hokkaido University, the scanners (i.e., 3 Tesla Magnetom Prisma, Siemens), number of channels in a coil, and imaging parameters were identical to each other.

### Preprocessing of MRI data

We performed preprocessing of the MRI data using the pipeline provided by fMRIPrep version 20.2.0 (Esteban et al., 2019). The preprocessing steps included slice-timing correction, motion correction, segmentation of T1-weighted structural images, coregistration, and normalization to Montreal Neurological Institute (MNI) space (for more details on the pipeline, see http://fmriprep.readthedocs.io/en/latest/workflows.html). The first five volumes of each run were discarded. We next applied spatial smoothing to the data with a 6-mm full-width at half-maximum (FWHM) Gaussian kernel using SPM 12 (Wellcome Trust Centre for Neuroimaging, London, UCL) on MATLAB. Spatial smoothing was not applied to the data for MVPA, since this might blur the fine-grained information contained in multivoxel activity (Mur et al., 2009).

We calculated frame-wise displacement (FD) based on the six realign parameters as an index of the amount of head motion for each run in each participant (Power et al., 2012). We excluded the runs in which the number of frames with FD > 0.5 mm was over 5% of all frames in a run from further analysis. As a result, the number of runs of five participants’ data became less than five; thus, we excluded these five participants from data analysis.

### General linear model analysis

We used a general linear model (GLM) analysis to explore activations or deactivations in brain regions after changing the level of accelerator sensitivity. One run included four conditions involving the sensitivity change: (1) change from low to high sensitivity, (2) change from high to low, (3) no change from low sensitivity, and (4) no change from high sensitivity. The following three periods of straight-section driving in each of the four conditions were modeled as separate 12-boxcar regressors that were convolved with a canonical hemodynamic response function: (1) a baseline period (period between the timings when the participant’s car passed the 100-m point of the straight section and when the preceding car passed the 400-m point), (2) an acceleration period (period between the timings when the preceding car passed the 400-m point of the straight section and when the participant’s car passed the 500-m point), (3) a target period (period between the timings when the participant’s car passed the 500-m point of the straight section and when the preceding car passed the 1000-m point). Note that we excluded the period for the first 100-m driving on the straight section from the baseline period to avoid contamination of the baseline period by the effect of curve driving. The six realign parameters were modeled as a regressor of no interest. We removed low-frequency noise using a high-pass filter with a cut-off period of 128 s. Serial correlations among scans were estimated with an autoregressive model implemented in SPM12.

To explore regions that were activated after the change in the level of accelerator sensitivity, we generated a contrast image using a fixed-effects model. We first generated contrasts of the target vs. baseline period in both the change and no-change conditions. We then subtracted the contrast in the no-change from that in the change condition:

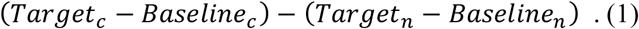

Here, *Target*_*c*_ and *Baseline*_*c*_ represent the target and baseline periods, respectively, of the change condition. *Target*_*n*_ and *Baseline*_*n*_ represent the target and baseline periods, respectively, of the no-change condition. Next, we explored regions that were deactivated after the sensitivity change by subtracting the contrasts in the opposite direction:

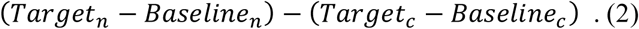

We also generated contrasts of the acceleration vs. baseline period using trials in both change and no-change conditions together in order to investigate brain regions that were activated or deactivated in response to acceleration of the preceding car. The contrast images of all participants were taken into the second-level group analysis using a random-effects model of a one-sample *t*-test. We adopted statistical inference with a threshold of *p* < 0.05 (family-wise error (FWE) corrected at cluster level with a cluster-forming threshold of *p* < 0.001). The anatomical localization was determined according to the automated anatomical labeling (AAL) atlas (Tzourio-Mazoyer et al., 2002).

### Multi-voxel pattern analysis

We performed MVPA to classify whether participants were driving the car with either high or low accelerator sensitivity from the fMRI voxel patterns (Haynes and Rees, 2005; Kamitani and Tong, 2005; Norman et al., 2006). We modeled the target period of each of the eight trials as a separate boxcar regressor that was convolved with a canonical hemodynamic response function. The GLM also included the six realign parameters as a regressor of no interest. As a result, eight independently estimated parameters (*β* values) per run were yielded for each voxel. We used the parameter estimates within a region of interest (ROI) as the input of a classifier. We used the linear support vector machine (SVM) implemented in LIVSVM (http://www.csie.ntu.edu.tw/~cjlin/libsvm/) with default parameters (a fixed regularization parameter C = 1). We targeted the anterior and posterior cerebellum as ROIs, which were anatomically defined with the AAL atlas and split into the left and right sides.

The SVM classifier was trained using the data in the no-change conditions (Fig. 2, left). We first aimed to examine whether the cerebellar activity exhibited distinct patterns that corresponded to the low- and high-sensitivity levels. We thus applied the trained classifier to the data in the no-change conditions (“test 1” in Fig. 2, right) and obtained a classification accuracy. Next, we aimed to examine whether the cerebellum switched its activity patterns according to the change in the level of accelerator sensitivity. Here, we applied the trained classifier to the data in the change conditions (“test 2” in Fig. 2, right). We performed a leave-one-run-out cross-validation in both “test 1” and “test 2” procedures to estimate classification accuracies as a way to prevent the differences among runs from becoming a confounding factor.

**Figure 2.**
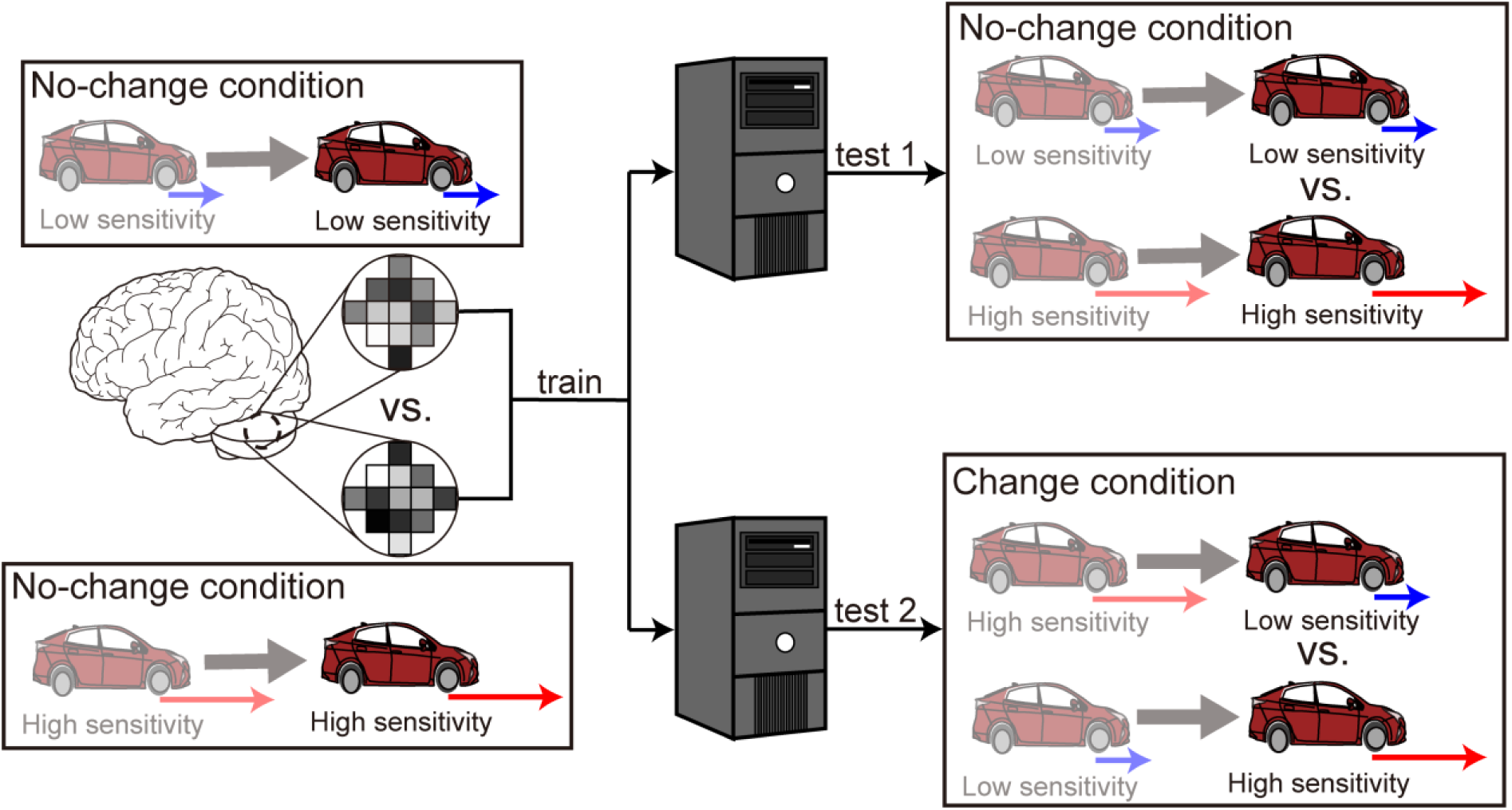
Multivoxel pattern analysis procedure. First, a classifier was trained to discriminate high from low sensitivity using the fMRI data in the no-change conditions. The trained classifier was applied separately to the data in the no-change conditions (denoted as “test 1”) and the data in the change conditions (denoted as “test 2”). The anterior and posterior parts of the cerebellum split into the left and right sides are selected as ROIs.

We statistically tested decoding accuracies by calculating z-scores using the following permutation procedure (Langfelder et al., 2011; Shibata et al., 2016; Ohata et al., 2020). We first randomly permuted the correspondence between the fMRI activity and the condition labels (i.e., high vs. low sensitivity) of the training dataset and then applied the classifier trained using the permuted data to the test dataset. We generated 1,000 surrogate classification accuracies by repeating the procedure 1,000 times. We calculated the *z*-scores of the original (without permutation) classification accuracies based on the empirical distribution of the 1,000 surrogate data for each participant separately. We tested the statistical significance of the *z*-scores using a two-sided *t*-test with a threshold of *p* < 0.05.

## Results

### Behavioral results

Figure 3A shows time courses of the inter-vehicle distance (distance between the preceding and participant’s driving cars) for a typical participant. The time courses were aligned at the possible timing of change in accelerator sensitivity (0 s: timing of passing the 500-m point of the straight section, see Fig. 1B). His car speeded up after the change in the level of accelerator sensitivity from low to high, resulting in the shortened inter-vehicle distance (Fig. 3A, left). By contrast, his car slowed down due to the sensitivity change from high to low, resulting in the lengthened distance (Fig. 3A, right). Once the inter-vehicle distance reached the maximum or minimum, the distance gradually returned to the target distance (25 m). Next, we subtracted the inter-vehicle distance in the no-change condition from that in the change condition at every sampling point. Figure 3B shows the difference averaged across participants. The difference in the inter-vehicle distance became largest at 9.6 s (averaged across participants, SD: 4.5 s) after the accelerator sensitivity changed from low to high and at 12.0 s (SD: 6.0) after the sensitivity changed from high to low.

**Figure 3.**
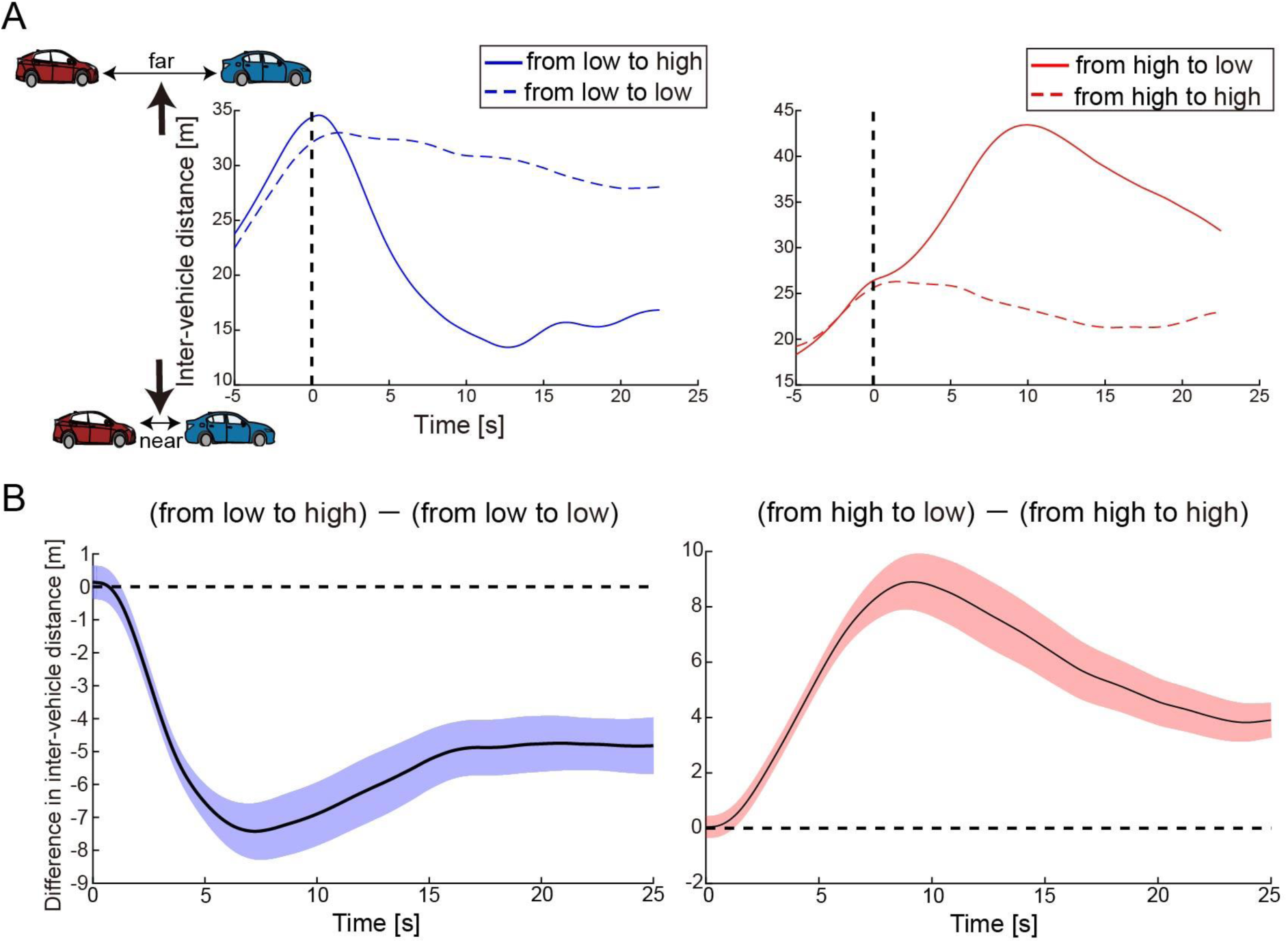
Behavior results. (A) Time courses of inter-vehicle distance for a typical participant. The solid blue and red lines denote time courses of the distance in the conditions when accelerator sensitivity changed from low to high and from high to low, respectively. By contrast, the dotted blue and red lines indicate time courses of the distance in the conditions where the sensitivity remained low or high, respectively. The distance was averaged across trials in each condition. The vertical dotted line denotes the timing at which his car passed the 500-m point of the straight section. In the change conditions, the level of accelerator sensitivity changed at this timing. (B) Time courses of difference in inter-vehicle distance between change and no-change conditions. The values were averaged across participants at every moment. Colored shaded areas indicate 95% confidence intervals.

### GLM results

We explored the brain regions in which the activities responded to the change in the level of accelerator sensitivity according to the contrast of activity represented in Equation (1). Figure 4A shows the regions activated after the sensitivity change (see Table 1). We first found significant activation in the left cerebellum, cerebellar lobule VI. We also found activation in the bilateral supramarginal gyrus and right middle frontal gyrus, which are portions of the inferior parietal lobe (IPL) and dorsolateral prefrontal cortex (DLPFC), respectively. The bilateral anterior insula and the thalamus, which are the components of the salience network (AI, Seeley et al., 2007; Menon and Uddin, 2010; Seeley, 2019), also activated after the sensitivity change. Next, we explored the deactivations in response to the sensitivity change (Fig. 4B and Table 1) according to the contrast represented in Equation (2). The posterior cingulate cortex (PCC), which is a component of the default mode network (DMN, Raichle et al., 2001; Buckner et al., 2008), and a part of the right cerebellum (VIIA CrusI) were significantly deactivated in response to the change in the level of accelerator sensitivity.

**Figure. 4.**
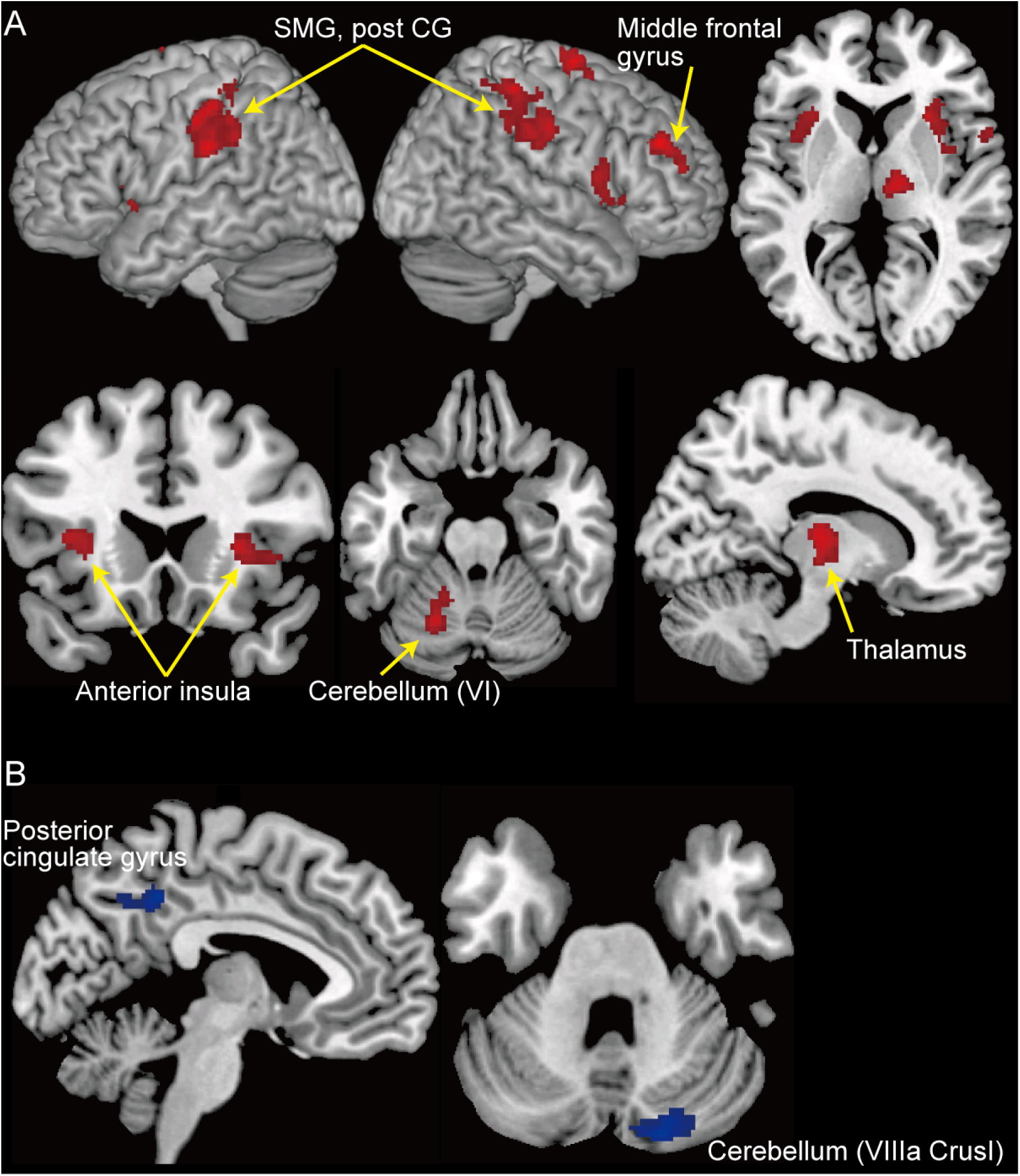
GLM results. (A) Clusters of activation (in red) that significantly increased after change in the level of accelerator sensitivity. (B) Clusters of activation (in blue) that significantly decreased after change in the level of accelerator sensitivity. A threshold at *p* < 0.05 (FWE-corrected at cluster level with a cluster-forming threshold of *p* < 0.001) was set for statistical testing. Post CG: post central gyrus, SMG: supramarginal gyrus.

**Table 1.** Summary of GLM results.

| Brain region | Side | Cluster size | *t*-value at peak | MNI coordinates<br>(peak voxel) |  |  |
| --- | --- | --- | --- | --- | --- | --- |
|  |  |  |  | x | y | z |
| Activate |  |  |  |  |  |  |
| 1. Thalamus | Right | 282 | 6.53 | 16 | -16 | 12 |
| 2. Supramarginal gyrus | Left | 647 | 6.06 | -66 | -26 | 28 |
| 3. Superior frontal gyrus | Right | 208 | 6.05 | 24 | -4 | 72 |
| 4. Middle frontal gyrus | Right | 156 | 6.00 | 36 | 40 | 32 |
| 5. Postcentral gyrus | Right | 624 | 5.91 | 38 | -34 | 58 |
| 6. Cerebellum (VI) | Left | 190 | 5.46 | -20 | -60 | -22 |
| 7. Insula | Right | 290 | 5.14 | 28 | 20 | 6 |
| 8. Insula | Left | 204 | 4.94 | -30 | 16 | 12 |
| 9. Opercular part of inferior frontal gyrus | Right | 144 | 4.71 | 54 | 10 | 14 |
| Deactivate |  |  |  |  |  |  |
| 1. Cerebellum (VIIa Crus1) | Right | 252 | 6.21 | 16 | -84 | -34 |
| 2. Posterior cingulate gyrus | Left | 158 | 5.00 | -2 | -44 | 42 |

A threshold at $p < 0.05$ (FWE-corrected at cluster level with a cluster-forming threshold of $p <$ 0.001) was set for statistical testing.

Next, we investigated brain regions activated by the acceleration of the preceding car. We generated the contrast between the acceleration and baseline periods from the data in both change and no-change conditions together. We found that some activated regions overlapped those found in Figure 4 (Supplementary Fig. 1). We were concerned that the activations in response to the sensitivity change were caused by contamination of the effect of the preceding car’s acceleration. To assess this concern, we generated the contrasts (acceleration vs. baseline periods) from the data in the change and no-change conditions separately. We compared the two contrasts but found no region showing a significant difference in the contrasts. Therefore, the activations/deactivations shown in Figure 4 were very likely involved in the change in the level of accelerator sensitivity.

### MVPA results

We first examined whether the cerebellar activity distinguished the two levels of accelerator sensitivity (high vs. low sensitivity) while participants were driving their car. We performed the classification analysis using the data only in the no-change condition (“test 1” in Fig. 2). As a result, we did not find any significant above-chance classification accuracy in any cerebellar ROI: 50.6 % (SD: 12.9 %) in the left anterior cerebellum (*t*(19) = -0.29, *p* = 0.77), 48.9 % (13.2 %) in the right anterior cerebellum (*t*(19) = -0.31, *p* = 0.77), 48.7 % (11.5 %) in the left posterior cerebellum (*t*(19) = -0.50, *p* = 0.62), and 52.5 % (12.0 %) in the right posterior cerebellum (*t*(19) = 0.99, *p* = 0.33). We next applied the classifier trained using the data in the no-change condition to those in the change condition (“test 2” in Fig. 2). The accuracy in no ROI was significantly higher than the chance level: 48.9 % (7.02 %) in the left anterior cerebellum (*t*(19) = -0.74, *p* = 0.47), 51.8 % (7.03 %) in the right anterior cerebellum (*t*(19) = -0.97, *p* = 0.34), 47.8 % (8.76 %) in the left posterior cerebellum (*t*(19) = -1.26, *p* = 0.22), and 51.5 % (8.92 %) in the right posterior cerebellum (*t*(19) = 0.83, *p* = 0.42). Accordingly, contrary to our expectation, the cerebellum did not maintain distinct activity patterns corresponding to the two levels of accelerator sensitivity or switch patterns in response to the sensitivity change.

## Discussion

The current fMRI study investigated the neural bases supporting driving behaviors adaptive to changes in the driving environment. We first found the activities in the sensorimotor area, including the cerebellum, the FPN, and AI regions, that increase in response to abrupt changes in the control property of a virtual car (Fig. 4A). By contrast, we found deactivation in a component of the DMN, the PCC, after the change (Fig. 4B). The GLM results suggest that the sensorimotor system is engaged in adaptive behaviors for car driving. Our results also highlight the role of the large-scale networks related to saliency, attention, and cognition in skillful driving.

Drivers are required to construct and switch multiple driving modes in response to changes in driving environments. Internal models may be the key mechanism underlying such adaptive driving behavior. Previous studies, in which participants controlled a computer mouse or joystick, demonstrated the cerebellar activity that reflects the acquisition of internal models (Imamizu et al., 2000; Imamizu et al., 2003; Seidler and Noll, 2008; Kim et al., 2015). The current driving study also reveals that the activity in the left cerebellar lobule VI significantly increases after the change in the level of accelerator sensitivity (Fig. 4A). Importantly, the activated area found in our study largely overlaps that involved in velocity control, not positional control, of a cursor manipulated with a computer mouse (Imamizu et al., 2003). This finding suggests the possibility that internal models for the same control property are allocated to the activity in the same cerebellar region regardless of the type of control device.

We also found activation in the DLPFC, IPL, and insula (Fig. 4A), all of which were reported to be associated with internal model switching (Imamizu et al., 2004). It has been suggested that the DLPFC and IPL contribute to internal model switching driven by the error between predicted and actual sensorimotor feedback (postdictive switching, Imamizu and Kawato, 2008). Forward internal models predict sensorimotor feedback from the efference copy of a motor command and are influential in the comparison between predicted and actual feedback (Wolpert et al., 1995; Miall and Wolpert, 1996; Blakemore et al., 2000). In our task, the level of accelerator sensitivity randomly changed in a trial-by-trial manner so that internal models could not be switched in advance of the sensitivity change (i.e., predictive switching). Thus, the model optimized for the environment before the sensitivity change addressed the new environment, resulting in a large prediction error. Our results suggest that such prediction error functioned as a signal of the change in the driving environment and prompted the brain to switch internal models postdictively.

Although the GLM results suggest the possibility of internal model switching, the MVPA results show neither distinction nor switching of cerebellar activity patterns during car driving. One possible reason for this unexpected result is that internal model switching is not necessarily an optimal strategy for every participant to address changes in the driving environment. Driving engages executive cognitive functions, such as attentional control, cognitive inhibition, and cognitive flexibility. Some participants might dominantly exert cognitive strategies in which such executive functions are employed rather than internal model switching. The driving simulator used in this study did not provide a perfectly realistic driving experience. The difference between real and simulated driving experiences might encourage participants to develop cognitive strategies uniquely optimal to the simulated environment. We found a large variance in the classification accuracies across participants, which might imply individual differences in their strategies.

We also found deactivation in the PCC in response to the change in the level of accelerator sensitivity (Fig. 4B). The PCC is a part of the DMN, a set of coordinated brain regions that shows more activation at rest than during cognitive demand tasks (Raichle et al., 2001; Buckner et al., 2008). The DMN regions respond to non-stimulus induced or internally oriented thoughts such as mind-wandering (Andrews-Hanna et al., 2010; Raichle, 2015), and thus the activation could interfere with task performance (Eichele et al., 2008; Hinds et al., 2013). The change in the driving environment possibly required participants to switch their cognitive modes of car-driving in addition to internal model switching. The deactivation in the PCC encouraged them to switch such cognitive modes quickly. The activities in the DMN regions are typically anticorrelated to those in the FPN regions (Fox et al., 2005; Fox et al., 2009). Previous studies suggest the crucial role of the SN in switching this balance between the FPN and DMN (Sridharan et al., 2008; Menon and Uddin, 2010; Uddin, 2015). Importantly, the current study found significant activation in the bilateral anterior insula and thalamus, parts of the SN (Fig. 4A). Our results imply the involvement of the large-scale networks (FPN, DMN, and SN) and their switching mechanism in adaptive driving behavior, which was a possible candidate of unique neural features for car driving.

As mentioned above, our driving simulator cannot represent a perfect driving experience. This issue constitutes the limitation of our current study. However, our task required participants exerting basic driving skills, such as speed control relative to a preceding car and changes in driving mode depending on the environment. No detailed examination of these skills has been carried out, to our knowledge, in previous functional neuroimaging studies. The best approach to overcoming this limitation is to develop a device that can measure brain activity, with comparable precision to an fMRI scanner, in a real car. Because the current technology cannot offer such devices, we translated driving behaviors into possible naturalistic behaviors inside an fMRI scanner while minimizing head movements. In our simulator, the accelerator and brake pedals were replaced by hand levers while the steering wheel was replaced by a knob dial. Such substitutions are common in the controllers of remote-control cars. Future studies are needed to improve both the driving simulator and neuroimaging devices in order to measure brain activities in more realistic driving experiences than that of the current study.

In sum, this study searched for the neural substrates supporting the adaptive behaviors of car driving. We found that the neural mechanisms studied in previous motor control studies, specifically internal model switching, are fundamental to adaptive driving behavior. Furthermore, this study also highlighted the involvement of the large-scale brain network in addressing cognitive demands for driving, suggesting a possible neural process unique to car driving.

## Data availability statement

The data for this study are available upon request to the corresponding authors.

## Ethics statement

The study was reviewed and approved by the Ethics Committee of the University of Tokyo. The participants provided their written informed consent to participate in this study.

## Author contributions

R.O. and H.I. designed the study; R.O. and K.O. collected the data; R.O. analyzed the data; R.O. and H.I. wrote the manuscript; K.O. reviewed and approved the final version of the manuscript.

## Acknowledgments

We are also grateful to Terumasa Endo, Satoshi Inoue, and Takeshi Hamaguchi (TOYOTA Motor Corporation) for their help in the experimental setup and insightful discussions about the results. We finally thank Shuhei Nishida and Yusuke Haruki for their assistance in collecting the data.

