## Supplementary Information for "Neural basis for adaptive motor behavior during car driving"

**Supplementary Figure**

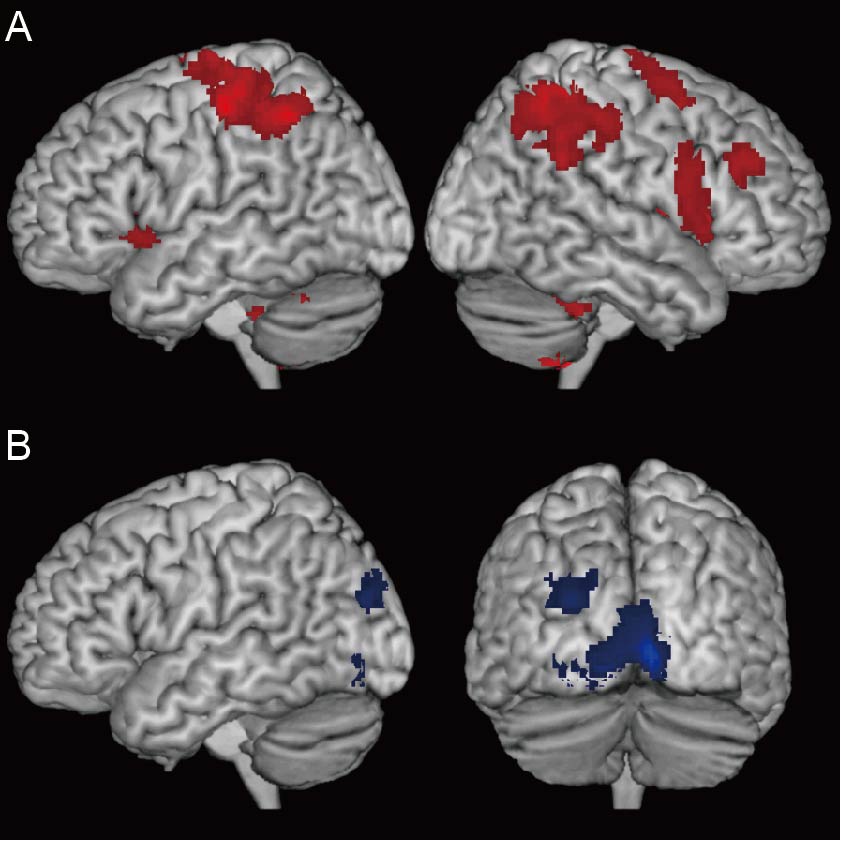

**Supplementary Figure 1**. Brain regions activated or deactivated in response to acceleration of preceding car. (A) Clusters of activation (in red) that significantly increased after the preceding car accelerated. (B) Clusters of activation (in blue) that significantly decreased after the preceding car accelerated. The threshold for statistical testing was set at *p* < 0.05 (FWE-corrected at cluster level with a cluster-forming threshold of *p* < 0.001).

**Supplementary Movie S1**. Virtual car driving in a straight section (related Figure 1B). The movie’s action is shown from the perspective of a driver (participant). The preceding blue car is seen in front of the participants’ driving car. The movie begins at the mid-point of the baseline period and finishes at the end of the target period (see Fig. 1B).
